# Hypoxia promotes a perinatal-like progenitor state in the adult murine epicardium

**DOI:** 10.1101/2021.09.16.460580

**Authors:** Angeliqua Sayed, Szimonetta Turoczi, Francisca Soares-da-Silva, Giovanna Marazzi, Jean-Sébastien Hulot, David Sassoon, Mariana Valente

**Affiliations:** Université de Paris, INSERM- U970, PARCC, F-75006 Paris, France; CIC1418 and DMU CARTE, AP-HP, Hôpital Européen Georges-Pompidou, F-75015, Paris, France; Lymphocytes and Immunity Unit, Immunology Department, Institut National de la Santé et de la Recherche Médicale U1223, Institut Pasteur, Paris, France

**Keywords:** Hypoxia priming, adult epicardial progenitors, in vitro epicardial system, multipotency stimulation, perinatal progenitor potential

## Abstract

The epicardium is a reservoir of progenitors that give rise to coronary vasculature and stroma during development and mediates cardiac vascular repair in lower vertebrates. However, its role as a source of progenitors in the adult mammalian heart remains unclear due to lack of clear lineage markers and single-cell culture systems to elucidate epicardial progeny cell fate. We found that *in vivo* exposure of mice to physiological hypoxia induced adult epicardial cells to re-enter the cell cycle and to express a subset of developmental genes. Multiplex transcriptional profiling revealed a lineage relationship between epicardial cells and smooth muscle, stromal, and endothelial fates, and that physiological hypoxia promoted an endothelial cell fate. *In vitro* analyses of purified epicardial cells showed that cell growth and subsequent differentiation is dependent upon hypoxia, and that resident epicardial cells retain progenitor identity in the adult mammalian heart with self-renewal and multilineage differentiation potential. These results point to a source of progenitor cells in the adult heart that can promote heart revascularization, providing an invaluable *in vitro* model for further studies.

## Introduction

The epicardium gives rise to epicardial-derived cells (EPDCs) located in the subepicardial layer that migrate into the myocardium to give rise to interstitial cells, perivascular stroma and smooth muscle cells during development and early postnatal life(Cai *et al*, 2008; Dettman *et al*, 1998; Gittenberger-de Groot *et al*, 1998; Mikawa & Gourdie, 1996; Perez-Pomares *et al*, 1997; Wessels *et al*, 2012; Zhou *et al*, 2008). The contribution of the epicardium to endothelial cells is controversial in mammals(Cai *et al.*, 2008; Duim *et al*, 2015; Merki *et al*, 2005; Perez-Pomares *et al*, 2002; Red-Horse *et al*, 2010; Tian *et al*, 2013; Zhou *et al.*, 2008) in part due to the small number of epicardial cells in the heart and a lack of suitable genetic cell fate models(Christoffels *et al*, 2009; Duim *et al.*, 2015; Duim *et al*, 2016; Rudat & Kispert, 2012). In addition, methodologies available to functionally study epicardial cells are limited to explant cultures that do not allow for definitive lineage and clonal analyses(Bax *et al*, 2019; Cao & Poss, 2016; Kim *et al*, 2012; Ramesh *et al*, 2016; Smart *et al*, 2007; Tran *et al*, 2016). Opposite to mammals, the multilineage capacity of the epicardium, including a contribution to endothelial cells, is well established in several non-mammalian vertebrate models where the adult epicardium gives rise directly to vasculature following injury(Dettman *et al.*, 1998; Manner, 1993; Merki *et al.*, 2005; Mikawa & Gourdie, 1996; Olivey *et al*, 2004; Poss *et al*, 2002).

In contrast to the adult, the neonatal heart is able to functionally recover following injury(Haubner *et al*, 2012; Porrello *et al*, 2011; Sampaio-Pinto *et al*, 2018). The perinatal heart is known to be hypoxic due to an immature vasculature structure(Nanka *et al*, 2006; Olivetti *et al*, 1980; Tian *et al.*, 2013; Tian *et al*, 2015; Tomanek, 1996) and other studies have shown that many progenitors in different tissues are present in hypoxic niches(Huang *et al*, 2018). These observations directly bear upon potential therapeutic avenues of discovery since it was reported recently that stepwise exposure of mice to hypoxia leads to a marked improvement in functional recovery of the heart following ischemia(Nakada *et al*, 2017). The degree of progenitor involvement in response to hypoxia in the adult remains unclear. The adult epicardium consists of progenitors and constitutes a physiological hypoxic niche due to the low capillary density and the constitutive expression of hypoxia inducible factor 1 alpha (HIF-1α)(Kocabas *et al*, 2012).

While these findings show that hypoxia and signaling effectors in a hypoxic response play a role in epicardial development and cardiac injury (Hesse *et al*, 2021a; Hesse *et al*, 2021b; Jing *et al*, 2016; Kocabas *et al.*, 2012; Tao *et al*, 2018), it remains unknown whether there is a stimulation (priming) of the adult epicardium after *in vivo* hypoxia exposure. In addition, how prospectively isolated single epicardial cells behave is unknown, since efforts to date have been hampered by the inability to maintain and expand these cells in culture. Consequently, the functional analysis of epicardial cells, at the single cell level, in respond to hypoxia has remained largely unaddressed. We therefore tested whether hypoxia regulates the epicardial progenitor competence. As a first step, we established a sorting strategy based upon known surface proteins together with a reporter mouse line for *Peg3/Pw1* (hereinafter referred to as *Pw1*). *Pw1* is expressed in a wide array of progenitor cells during development and in the adult, and is linked to the stem cell capacity of self-renewal as well as differentiation into specific tissue lineages(Besson *et al*, 2011; Kuroiwa *et al*, 1996; Mitchell *et al*, 2010). We demonstrated previously that PW1 is expressed in a subset of stromal cells in the adult heart, which generates pro-fibrotic cells in response to injury(Bouvet *et al*, 2020; Yaniz-Galende *et al*, 2017). However, PW1 expression had not been examined during early postnatal development when rapid heart growth and differentiation takes place. We show here that PW1 is expressed in the Gp38^+^ epicardium (Mahtab *et al*, 2008; Smart *et al*, 2011; Valente *et al*, 2019) from development throughout adult life, consistent with progenitor capacity for this compartment(Cai *et al.*, 2008; Dettman *et al.*, 1998; Gittenberger-de Groot *et al.*, 1998; Mikawa & Gourdie, 1996; Perez-Pomares *et al.*, 1997; Wessels *et al.*, 2012; Zhou *et al.*, 2008). We also show that hypoxia induces an increase in the number of PW1 expressing cells in the Gp38^+^ epicardium and PDGFRα^+^ subepicardium. Multiplex transcriptional profiling revealed that Gp38^+^PW1^+^ epicardial cells upregulate endothelial lineage-associated genes in response to hypoxia similar to what is found in the epicardium at birth. Lastly, while previous studies relied on explants cultures, we show here that freshly purified Gp38^+^PW1^+^ epicardial cells display robust clonogenicity, self-renewal, and multipotency uniquely under hypoxic conditions, whereas they do not grow under normoxic conditions. Adult Gp38^+^PW1^+^ epicardial cells from physiological hypoxia-primed mice displayed a marked increase in their competence to grow and differentiate *in vitro* similar to what is observed with the neonatal epicardial cells. Taken together, our data support the hypothesis that the adult mammalian epicardium is a niche for resident Gp38^+^PW1^+^ progenitors and that exposure to hypoxia promotes robust progenitor activity similar to neonatal epicardium, including the activation of endothelial cell fate.

## Results

### Chronic physiological hypoxia exposure in the adult heart activates a developmental profile in the epicardial and subepicardial layers

Previous studies have shown that the epicardium is hypoxic (Kocabas *et al.*, 2012) and contains multipotent progenitors (Acharya *et al*, 2012; Smart *et al.*, 2011; Wessels & Pérez-Pomares, 2004; Winter & Gittenberger-de Groot, 2007). While there are conflicting reports regarding epicardial progenitor potential in the adult mammalian heart, it is well established that the epicardium has a pronounced progenitor capacity during mammalian development, corresponding to a hypoxic state in both the epicardium and underlying cell layers (Guimaraes-Camboa *et al*, 2015; Sugishita *et al*, 2004; Tomanek *et al*, 2003; Wikenheiser *et al*, 2006). This raised the possibility that hypoxia directly regulates epicardial progenitors and is required for progenitor function. To explore the role of hypoxia, we placed adult mice into a hypoxic chamber at 10% O_2_ for 2 weeks (Figure 1A), which is well tolerated and sufficient to induce overt physiological changes, including cardiac enlargement coupled with a reduced overall body weight, as reported by others(Nakada *et al.*, 2017) (Figure I-A-B in the Supplement). We used pimonidazole as a surrogate marker for hypoxic cells(Krohn *et al*, 2008; Nunn *et al*, 1995). Pimonidazole staining in the epicardium and subepicardium as well as a stronger staining in the myocardial interstitium confirmed that the experimental conditions used were sufficient to induce generalized hypoxia in the heart (Figure 1B-C) and we noted that pimonidazole labeling intensity was similar in to that detected at P0 (Figure 1C).

**Figure 1.**
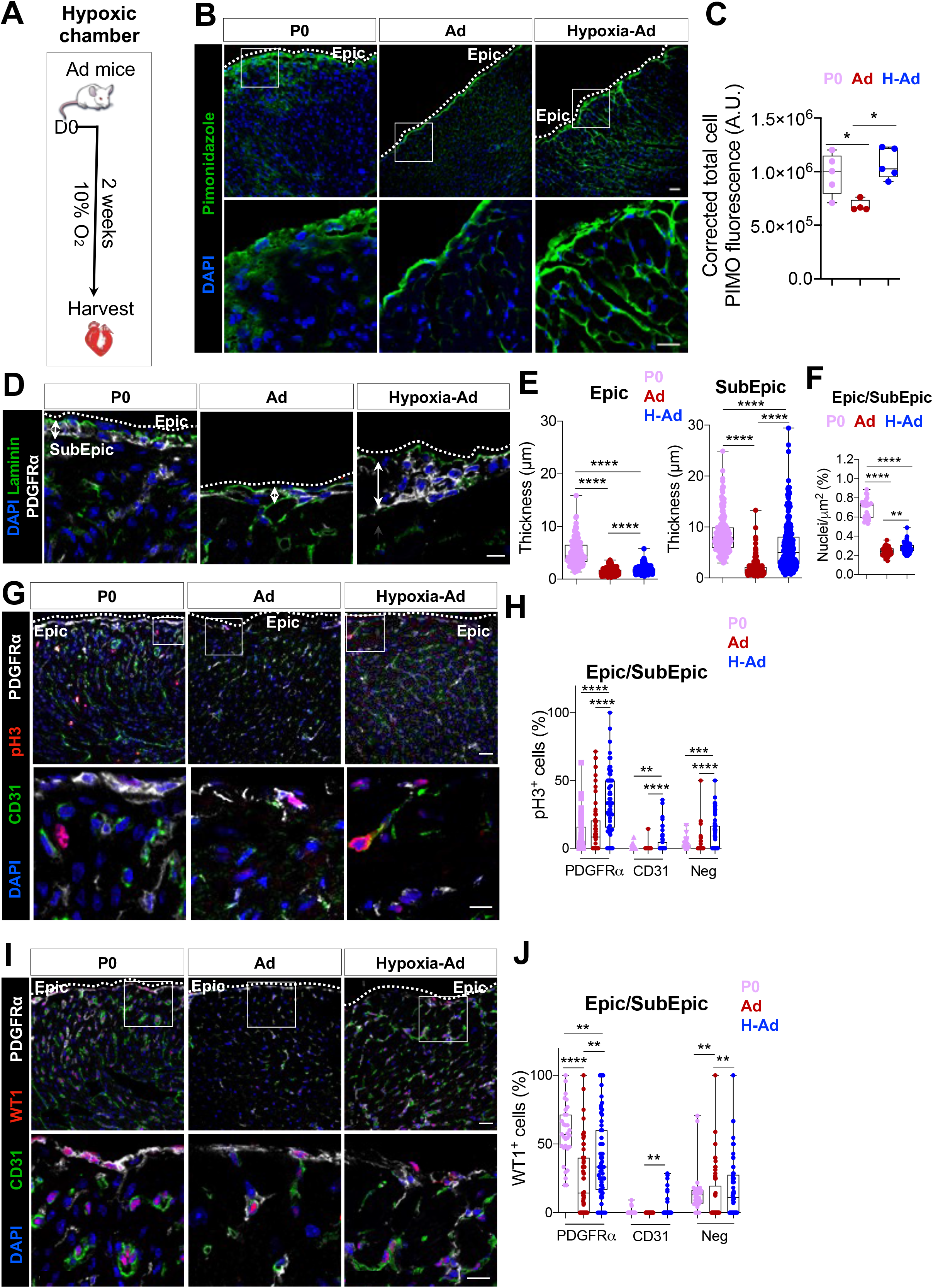
Two-weeks hypoxia exposure induces the reactivation of an epicardium-specific developmental growth program including proliferation and WT1 expression. A. Schema of chronic hypoxia induction (10% O_2_) in the adult mice. B. Hypoxic regions stain with Pimonidazole in P0, normoxic and hypoxic adult ventricles. Scale bar: 20μm (top panels), scale bar: 10μm (bottom panels). C. Pimonidazole expression increases under hypoxia exposure (n=3). D. Epicardial and subepicardial layer determined by PDGFRα and Laminin staining in P0, normoxic and hypoxic adult ventricles. Scale bar: 10μm. E. Increase of epicardial and subepicardial thickness in hypoxic mice (n=3, 20 images per heart, 5 measures per image and per layer). F. Higher number of nuclei in hypoxic mice (n=3, 20 images per heart). G. Proliferative PDGFRα^+^ and CD31^+^ cells determined with pH3, in normoxic and hypoxic adult ventricles. Scale bar: 20μm (top panels), 10μm (bottom panels). H. Overall increase of proliferative cell in hypoxic mice (n=3, 20 images per heart and per region). I. Immunostaining of WT1, a marker of epicardial activation. Scale bar: 20μm (top panels), 10μm (bottom panel). J. Increase of WT1 expression particularly in the epicardial, subepicardial and endothelial cells in hypoxic mice (n=3, 20 images per heart and per region). All nuclei were counterstained with DAPI. Dashed line delineates the epicardial (Epic) layer. Values are normalized by the total number of nuclei per layer and per field and all values are represented in percentage. The line in the box plot represents the median. Statistical significance was determined either by Mann-Whitney test or by one-way ANOVA (Kruskal-Wallis test) with uncorrected Dunn’s test, for 2 groups or 3 groups comparison, respectively. \*\*\*\**p*<0.0001, \*\*\**p*<0.001, \*\**p*<0.01. P0: postnatal day 0, Ad: Adult, H: Hypoxia, Epic: Epicardium, SubEpic: Subepicardium, Neg: Negative Fraction.

Histological analyses revealed a striking increase in the thickness of the epicardial and subepicardial layers in response to hypoxia (Figure 1D-E) coupled with an increase in nuclei number (Figure 1F), as observed during development (Figure I-D in the Supplement). Consistent with these observations, we detected a marked increase in cell proliferation in the subepicardial cells (PDGFRα^+^, Figure 1G-H) in response to hypoxia using the cell proliferation marker phosphorylated histone H3 (pH3), suggesting that cell proliferation contributes to the increased thickness following hypoxia in the subepicardial layer (Figure 1D-F). While proliferative cells are present at low levels in the adult heart(Eschenhagen *et al*, 2017; Soonpaa *et al*, 1996) our results show that chronic exposure to hypoxia activates epicardial and subepicardial cell proliferation. Previous studies have shown that proliferation of epicardial/subepicardial cells is a feature of embryonic development(Cai *et al.*, 2008; Dettman *et al.*, 1998; Gittenberger-de Groot *et al.*, 1998; Wessels & Pérez-Pomares, 2004; Zhou *et al.*, 2008), therefore we tested whether other epicardial developmental markers were induced concomitant with cell proliferation. One such marker, WT1, is expressed in the epicardial and subepicardial compartments during embryonic heart development (Cossette & Misra, 2011; Duim *et al.*, 2016; Manner *et al*, 2005; Zhou *et al.*, 2008). We observed a marked induction of WT1 expression in the epicardium and subepicardium (Figure 1I-J). Taken together, our results show that physiological hypoxia activates both cell proliferation and WT1 expression in the adult that are hallmarks of early epicardial/pericardial development.

### PW1 is expressed at high levels in the epicardium and subepicardium throughout life

In order to isolate epicardial and subepicardial cells to assess their progenitor potential profile, we established a precise gating strategy to purify the different cardiac populations. We used antibodies to multiple cell surface proteins combined with a reporter mouse model for the expression of the adult progenitor cell marker *Peg3/Pw1* (hereinafter referred to as *Pw1*)(Besson *et al.*, 2011; Kuroiwa *et al.*, 1996; Mitchell *et al.*, 2010). Endothelial and stromal cells were defined by the expression of platelet endothelial cell adhesion molecule 1 (PECAM-1^+^ or CD31^+^) and platelet-derived growth factor receptor alpha (PDGFRα or CD140a), respectively. Hematopoietic cells were excluded by the combination of CD45, CD11b and Ter119 antibodies (Figure IIA in the Supplement). To define epicardial cells, we used an antibody to the surface glycoprotein 38 (Gp38 or podoplanin, Figure IIA in the Supplement), which has been shown to be expressed in the epicardial layer during development(Mahtab *et al.*, 2008; Valente *et al.*, 2019) as well as in the adult(Smart *et al.*, 2011; Valente *et al.*, 2019). While Gp38 is expressed in other cardiac cells population (hematopoietic, endothelial and stromal cells)(Hou *et al*, 2010; Stellato *et al*, 2019; Weninger *et al*, 1999), we excluded these cell lineages in our sorting strategy (Figure 2A, Figure IIA-B in the Supplement). We found ~70% of the Gp38-expressing cells co-expressed the *Pw1* reporter gene at P0 and the *Pw1* reporter gene co-expression was maintained in the adult (≈ 50%, Figure 2B). We observed that PDGFRα^+^ cells (subepicardial/stromal cells) expressed the *Pw1* reporter gene at all stages at high levels (≈ 80% and 60%, respectively, Figure 2B), whereas CD31^+^ cells (endothelial cells) underwent a sharp decline in PW1 expression during postnatal life (P0 ≈ 45% vs. adult ≈ 10%, Figure 2B). Flow cytometry analyses revealed that while the overall frequency of PW1^+^ cells increases in response to hypoxia (Figure IIIA-B in the Supplement), this increase is limited to the Gp38^+^ epicardial and CD31^+^ endothelial cells, whereas stromal cells do not show a significant change (Figure 2B). Immunostaining confirmed PW1 protein expression in the epicardium and subepicardium and showed that PW1^+^ cells co-expressed Gp38 in the epicardium and PDGFRα in the subepicardium (Figure 2C-D). Although an overall ~4-fold decrease of PW1 expression was observed between P0 and adult in the epicardium (Gp38^+^) and subepicardium (PDGFRα^+^), PW1 expression remained markedly higher in these layers (Figure 2E). These results also reveal that subepicardial PDGFRα^+^ cells respond to hypoxia through an increase in PW1 expression, however the stromal PW1^+^PDGFRα^+^ compartment present in the cardiac interstitium do not show a significant response as measured by flow cytometry (Figure IIIC in the Supplement). We propose that the marked increase in PW1 expression in PDGFRα^+^ subepicardial cells is not observed using flow cytometry since the PDGFRα^+^ subepicardial cells constitute a minor population as compared to PDGFRα^+^ stromal cells which cannot be discriminated between the two compartments using the available cell surface markers.

**Figure 2:**
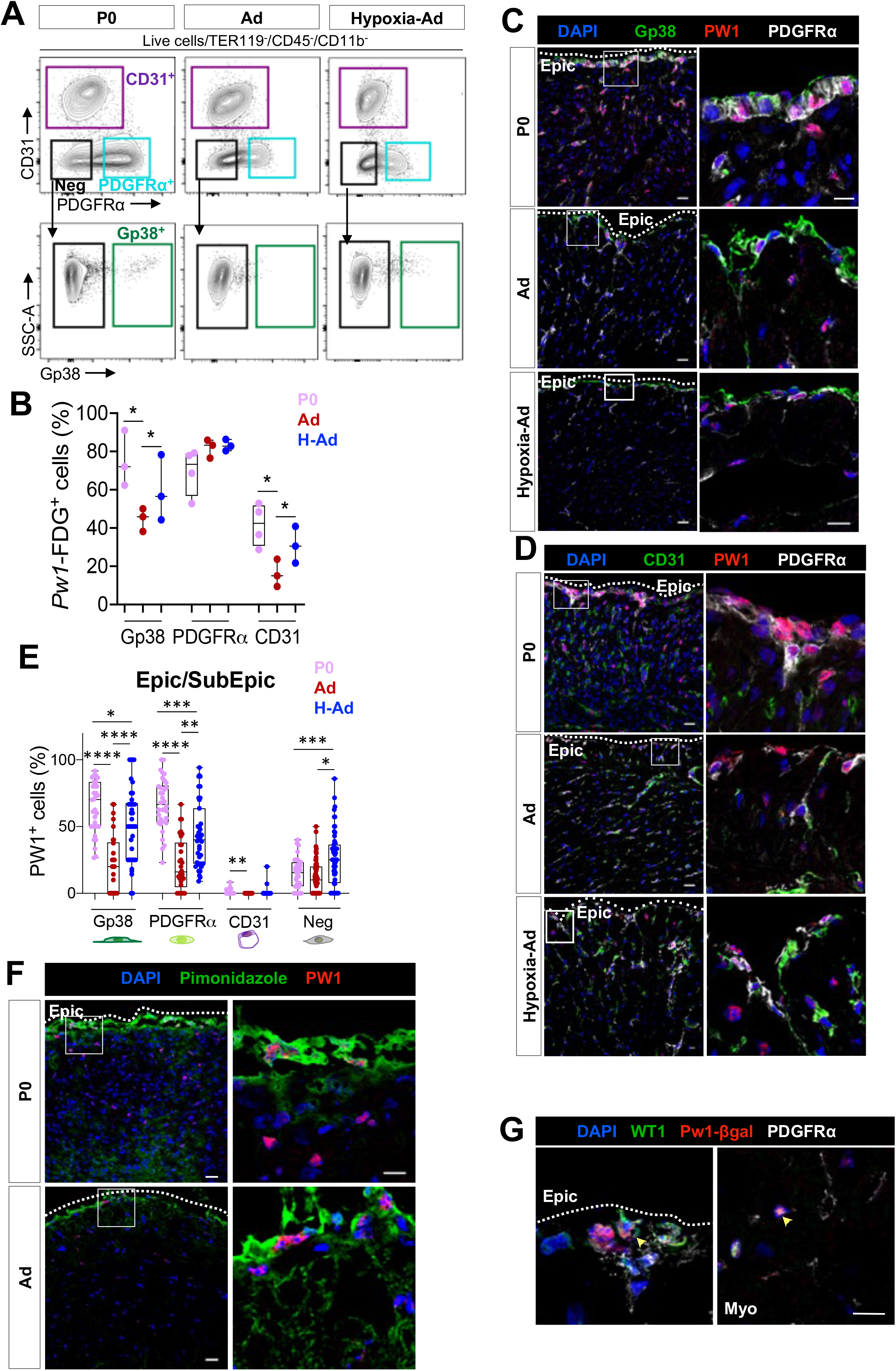
Adult mice exposed to chronic hypoxia (10%) upregulate PW1 expression in the activated epicardium and subepicardium. A. Flow cytometry profiles and gating strategy of P0, Ad, and Ad-Hypoxia cardiac cell suspensions stained with CD31, PDGFRα and Gp38. B. *Pw1*-FDG^+^ cells increase particularly in the endothelial and epicardial cell compartment after hypoxia induction in adult mice (n=3). C. Co-staining of PW1, Gp38 and PDGFRα in hypoxic mice (left panel). Scale bar: 20μm (upper), 10μm (bottom). D. Co-staining of PW1, CD31 and PDGFRα in hypoxic adult ventricles. Scale bar: 10μm. E. Increase of PW1 expression in the epicardium (Gp38^+^) and subepicardium (PDGFRα^+^ and negative cell fraction) in hypoxic adult ventricle (n=3, 20 images per heart and per region). F. Co-expression of PW1 with pimonidazole in the epicardium and subepicardium in P0 and adult ventricles. Scale bar: 20μm (left panels), 10 μm (right panels). G. Co-staining of Pw1-β-gal with WT1 and PDGFRα in hypoxic adult mice. Scale bar: 10μm. All nuclei were counterstained with DAPI. Values are normalized by the total number of nuclei per layer and per fields and all values are represented in percentage. Dashed line delineates the epicardial (Epic) layer. The line in the box plot represents the median. Statistical significance was determined by one-way ANOVA (Kruskal-Wallis test) with uncorrected Dunn’s test. \*\*\*\**p*<0.0001, \*\*\**p*<0.001, \*\**p*<0.01, \**p*<0.05. Epic: Epicardium, SubEpic: SubEpicardium, Myo: Myocardium, P0: postnatal day 0, Ad: Adult, H: Hypoxia, *Pw1*-FDG: *Pw1*-associated 5-Dodecanoylaminofluorescein Di-β-D-Galactopyranoside, PIMO: Pimonidazole.

We and others have shown that *Pw1* participates in multiple cell stress pathways(Feng *et al*, 2008; Kohda *et al*, 2001; Relaix *et al*, 2000; Relaix *et al*, 1998; Schwarzkopf *et al*, 2006) and regulates glucose metabolism(Correra *et al*, 2018). We therefore explored whether PW1 expression is regulated by hypoxia that is known to trigger cell stress and regulate glucose metabolism(Giaccia *et al*, 2004). We confirmed that hypoxic cells are restricted to the epicardial and subepicardial layers in the adult (Figure 1B) and express PW1 (Figure 2F). Moreover, the developmental epicardial transcription factor WT1 was co-expressed with the *Pw1*-reporter gene in the epicardium (Gp38^+^), subepicardium (PDGFRα^+^, Figure 2G and Figure IIID in the Supplement), indicating that the activated epicardial and subepicardial cells also co-expressed Pw1. Taken together, our data show that PW1 expression decreases during postnatal life in the ventricles, but levels remain elevated in the epicardial and subepicardial compartments. This observation is particularly relevant since epicardial and subepicardial layers have been shown to be a reservoir of cardiovascular progenitors during development(Cai *et al.*, 2008; Dettman *et al.*, 1998; Gittenberger-de Groot *et al.*, 1998; Wessels & Pérez-Pomares, 2004; Zhou *et al.*, 2008) and that PW1 is expressed by multiple progenitor cell types in the adult(Besson *et al.*, 2011; Kuroiwa *et al.*, 1996; Mitchell *et al.*, 2010).

### Epicardial cells express a progenitor/multipotent transcriptional profile in response to hypoxia

To better characterize the epicardial response to hypoxia and define how the adult epicardium compares to the perinatal state, we designed a multiplex qPCR panel of known genes corresponding to the major proposed epicardial lineages as well as to genes that respond to oxygen metabolism in freshly sorted cardiac populations (Table I in the Supplement). Unsupervised hierarchical clustering (Figure IV-A in the Supplement) and UMAP1 vs. UMAP2 (Uniform Manifold Approximation and Projection, Figure IV-B in the Supplement) at population level segregated the main cardiac populations in: CD31^+^ endothelial cells irrespective of the time-point or hypoxia exposure (cluster I); developing Gp38^+^ epicardial cells (E17.5 and P0, cluster II); the majority of PDGFRα^+^ cells assembled in two adjacent clusters (clusters III and IV), which differ from each other due to high expression of *Cdh5* and lower levels of *Col1a1*, *Tcf21* and *Postn* in the cluster of hypoxia-primed adult PDGFRα^+^ and Gp38^+^ epicardial cells (cluster IV, Figure IV in the Supplement); cluster V is heterogeneous and is composed of adult Gp38^+^ epicardial cells (both normoxic and hypoxic) together with hypoxia-primed PDGFRα^+^ and CD31^+^ cells (Figure IV in the Supplement). While these results validate our approach to discriminate the main cardiac populations, they also reveal clustering of Gp38+ cells with other cell types, reflecting heterogeneity of epicardial cells and the transcriptional profile overlap of hypoxia exposed epicardial, stromal, and endothelial cells (Figure IV in the Supplement).

We analyzed 382 cells at the single cell level, and further confirmed our results observed at the population level (Figure IV in the Supplement) by unsupervised hierarchical clustering (heat-map, Figure V-A in the Supplement) and tSNE projection (t-Distributed Stochastic Neighbor Embedding, Figure V-B in the Supplement). Freshly sorted Gp38^+^ epicardial cells are highly heterogeneous and were split in the four obtained clusters (Figure V in the Supplement). Further single cell transcriptional analysis of the Gp38^+^ epicardial population (168 cells) revealed unique transcriptional signatures of the different epicardial subsets (Figure 3A). Epicardial-associated genes (*Wt1*, *Gpm6a*, *Bnc1* and *Tbx18*) are highly expressed during development (P0 cluster I, III and IV) and after injury(Cossette & Misra, 2011; Duim *et al.*, 2016; Manner *et al.*, 2005; Zhou *et al.*, 2008). We observe a re-activation of these genes after hypoxia exposure in a fraction of cells at the transcriptional level (cluster III, Figure 3A), as well as at the protein level (Figure 1I-J). Cluster I encompass a compartment of P0 epicardial cells expressing the epicardial specific genes together with *Kdr*, *Flt1*, *Pdgfb*, *Nos3*, *Cdh5* and *Acta2*. Co-expression in the same cell of epicardial, endothelial and *Acta2* is compatible with an epicardial commitment to the endothelial lineage. Cluster II contains the majority of adult normoxic and hypoxic Gp38^+^ epicardial cells expressing stromal associated genes (*Col3a1*, *Tmsb4x*, *Col1a1*, *Tcf21* and *Tbx20*), confirming their lineage relationship as shown by others(Cai *et al.*, 2008; Dettman *et al.*, 1998; Gittenberger-de Groot *et al.*, 1998; Wessels & Pérez-Pomares, 2004; Zhou *et al.*, 2008). Cluster III and IV show high levels of epicardial specific genes (*Wt1*, *Gpm6a*, *Bnc1* and *Tbx18*) indicative of the most immature epicardial profile. Cluster III differs from cluster IV due to the presence of hypoxia-primed epicardial cells and the additional expression of *Cldn5*. This cluster also diverges from cluster II, where the majority of the hypoxia-primed epicardial cells are, due to the reduction of the fibrotic genes (*Tcf21, tbx2o, Tgfb1* and *Col1a1*, Figure 3A). This result reveals the acquisition of endothelial lineage transcript in the adult epicardial cells only following hypoxia exposure. The projection of the transcriptional data as UMAP1 vs. UMAP2 (Uniform Manifold Approximation and Projection, Figure 3B) show similarly to the heatmap clustering (Figure 3A), with 4 clusters of epicardial cells. The UMAP analysis highlights the similarities between cluster I (P0 epicardial cells) and cluster III (P0 and hypoxia-primed epicardial cells) with the differential expression of genes associated to the endothelial lineage (Figure 3B).

**Figure 3.**
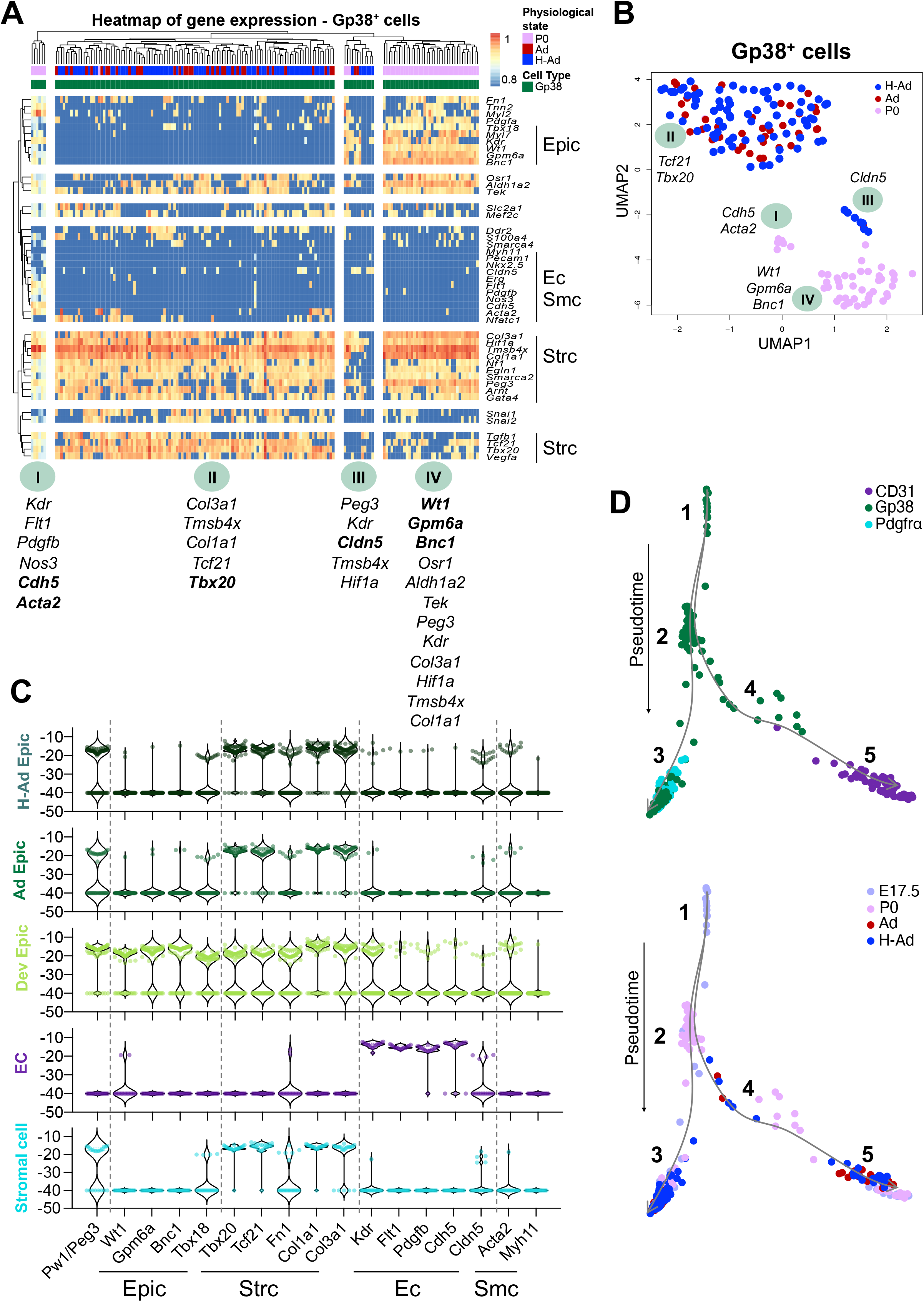
Transcriptional profiling demonstrates the activation of the epicardial developmental growth program in response to chronic hypoxia and a lineage overlap with PDGFRα^+^ and CD31^+^ cells. A. Multiplex qPCR heat-map at single cell level reveals heterogeneity of the Gp38^+^ epicardial compartment from P0, Ad (adult) and H-Ad (hypoxic adult). Each column represents one cell with the corresponding color-code according cell type and physiological state. Hierarchical clustering reveals 4 major group of cells labeled I, II, III, IV with the differentially expressed genes listed below. A total of 168 cells were analyzed. Gene expression was normalized to *Gapdh*, and unsupervised hierarchical clustering was performed. B. tSNE analysis of multiplex qPCR from 168 epicardial cells (P0, Ad and H-Ad). Genes displayed are the most differential expressed genes corresponding to each cluster and are color-coded according to the to the physiological state. C. Violin plot of genes corresponding to the major epicardial derived lineages (Epic: epicardium, Sc: Stromal cells, Ec: endothelial cells, Smc: smooth muscle cells) of *Pw1/Peg3* and of metabolic pathways from a total of 186 cells analyzed. D. Diffusion map (pseudo-time analysis) of Sc, Ec, and Smc populations based on single cell gene expression. Color-code according to the cell type (top panel) and according to the physiological state (bottom panel). E17.5: embryonic day 17.5, P0: postnatal day 0, Ad: Adult, H: Hypoxia, Epic: epicardium, Ec: endothelial cell, Smc: smooth muscle cell, Strc: stromal cells.

Based on single cell data, gene expression profiles corresponding to the major epicardial derived lineages, i.e. epicardial, stromal, endothelial and smooth muscle cells were analyzed (Figure 3C). While the developing epicardium shows a mixed profile with cells expressing genes characteristic of all lineages (epicardial, stromal, endothelial and smooth muscle cells), the adult epicardial layer only expresses epicardium and stromal cell transcripts. In contrast, hypoxia-primed adult epicardium up-regulates the expression of endothelial and smooth muscle-associated genes in a subset of cells comparable to the developing epicardium profile (Figure 3C).

While not definitive, the determination of lineage trajectories can be proposed based upon multiplex single-cell experimental tools and diffusion maps (or pseudotemporal ordering) as shown by others(Haghverdi *et al*, 2016). We therefore generated diffusion maps for the 3 populations examined here. Similar to our observations with unsupervised clustering and tSNE analyses (Figure IV and V-B in the Supplement), we found that Gp38^+^ epicardial cells were more dispersed, whereas PDGFRα^+^ stromal and CD31^+^ endothelial cells were found closer together (Figure 3D). The resulting trajectory initiates with the most immature phenotype (Gp38^+^ cells, root, labeled 1) and splits into two distinct cell fates, the stromal cell (labeled 3) and endothelial cell (labeled 5). Developing epicardial cells (E17.5) are positioned at the root of the diffusion map (Figure 3D, labeled 1), followed by P0 epicardial cells (Figure 3D, labeled 2) and a split node into two branches: one of PDGFRα^+^ cells (Figure 3D, labeled 3); and another of CD31^+^ endothelial cells (Figure 3D, labeled 5). We noted a large overlap between the PDGFRα^+^ cell branch and the Gp38^+^ epicardial cells consistent with a cell fate lineage relationship. Endothelial cell branch shows less overlap and is more clearly segregated from Gp38^+^ cells. However, we noted a fraction of hypoxia-primed adult Gp38^+^ epicardial cells that cluster together with P0 epicardial cells (Figure 3D, labeled 4) alongside the pathway of the endothelial cells fate. This result indicates that epicardial cells are distinct from the endothelial compartment. Overall, Gp38^+^ epicardial cells display a heterogeneous transcriptional profile for the genes analyzed consistent with the proposal that this compartment consists of progenitor cells that can give rise to multiple lineages. As expected, E17.5 and P0 derived epicardial cells have the most immature transcriptomes. A significant fraction of epicardial cells, irrespective of the stage or conditions from which they were isolated largely overlap with the PDGFRα^+^ population which is in agreement with previous demonstrations of a lineage relationship between the epicardium and stromal/fibroblast cells(Acharya *et al.*, 2012; Ali *et al*, 2014; Cai *et al.*, 2008; Dettman *et al.*, 1998; Gittenberger-de Groot *et al.*, 1998; Grieskamp *et al*, 2011; Katz *et al*, 2012; Mikawa & Gourdie, 1996; Perez-Pomares *et al.*, 1997; Wessels *et al.*, 2012; Zhou *et al*, 2011; Zhou *et al*, 2012a; Zhou *et al.*, 2008). In addition, we observed that a distinct subset of P0 and hypoxia-primed epicardial cells cluster with the endothelial population, highlighting a potential contribution to the endothelium. These data suggest that the adult epicardium is activated to give rise to a population of cells with an endothelial cell fate potential in response to hypoxia, which has been proposed by others to occur during cardiac development(Limana *et al*, 2007; Wessels *et al.*, 2012) as well as in non-mammalian vertebrates throughout life(Manner, 1993; Merki *et al.*, 2005; Olivey *et al.*, 2004; Poss *et al.*, 2002).

### The epicardial *in vitro* stem cell potential is dependent upon hypoxia

The cell fate potential of the epicardium can only be defined using reliable lineage markers which are presently unavailable. This limitation is further compounded by the currently available *in vitro* assays to study epicardial cells biology that are based on tissue explants and migratory potential of epicardial-derived cells (Bax *et al.*, 2019; Cao & Poss, 2016; Kim *et al.*, 2012; Ramesh *et al.*, 2016; Smart *et al.*, 2007; Tran *et al.*, 2016). Tissue explant assays do not allow for the determination of the main stem cell properties (i.e. self-renewal, clonogenicity and multipotency) and bias the study to cells able to migrate out of the explant. Our data reveals that epicardial cells are activated in response to hypoxia, raising the possibility that these cells require a hypoxic environment. Using our sorting strategy (Figure IIA in the Supplement), we prospectively isolated and freshly cultured Gp38^+^PW1^+^ epicardial cells from fetal (E17.5), newborn (P0), and adult hearts both in normoxic (21% O_2_) and in hypoxic (1% O_2_) conditions (Figure 4A). We found that Gp38^+^PW1^+^ epicardial cell growth was completely dependent upon hypoxic conditions regardless of their developmental state, whereas these cells did not expand in normoxic conditions and only few cells were present following several days in culture (Figure 4A-B). We noted that adult-derived epicardial cells displayed a significant decrease in the ability to grow in culture (Figure 4B). Because the ability of adult epicardial cells to grow in culture is decreased and the frequency of epicardial cells recovered from an adult heart is substantial smaller, we characterized the *in vitro* epicardial colonies potential using the developing epicardial Gp38^+^ cells (E17.5 and P0). Low-density plating assays showed that Gp38^+^PW1^+^ epicardial cells form colonies with an epithelial-like honeycomb morphology (Figure 4B-C). The developing (E17.5/P0) colonies displayed complete overlap of Gp38 and PW1 expression (Figure 4C), showing that hypoxia maintains their original profile (Figure 4A). Cell proliferation was assessed by Ki67 and pH3 expression and showed that ~95% cells (Gp38^+^/Ki67^+^) were proliferative of which 5% were actively dividing (Gp38^+^/pH3^+^, Figure 4D and Figure VII in the Supplement). This profile was observed at D4 or D8 in culture (Figure VI in the Supplement), confirming that colony growth is due to proliferation of Gp38^+^ cells. Additionally, colonies shared a similar average size when derived from E17.5 and P0 (Figure 4D, right graph) revealing a similar growth potential in cells derived from early stages. We next characterized the colonies for each relevant epicardial lineage following 8 days in culture. We observed that PDGFRα was expressed in a subset of the colony (Figure 4E and Figure VII in the Supplement) and its expression is acquired by D4 (Figure VI in the Supplement). This result is consistent with previous observations that PDGFRα is expressed in epicardial-derived cells found in the subepicardial zone (Figure 2C-D). The Gp38^+^PW1^+^ colonies also showed expression of the smooth the muscle cell markers αSMA and SM22α (Figure 4F and Figure VII in the Supplement). The majority of cells within the colony expressed both proteins, however the co-expression of αSMA was associated with a smooth muscle-like cell morphology. We detected that colonies typically consisted of a centrally located core of cells that were round and compact that expressed high levels of Gp38^+^ (Figure 4F, inset 1). This core was typically surrounded by an outer layer of differentiating αSMA^+^ cells with a lower nucleus/cytoplasm ratio and lower levels of Gp38 expression (Figure 4F, inset 2). Furthermore, we observed a widespread expression of Flk1, an early endothelial commitment marker (Figure 4G and Figure VII in the Supplement). While Flk1 expression confirms that the Gp38^+^ cells can give rise to the endothelial lineage, we did not observe CD31 expression, which likely reflects that our culture conditions, while suitable for cell expansion, are not optimized for full endothelial differentiation. We further confirmed the multipotential ability of the Gp38^+^Pw1^+^ cells to differentiate into smooth muscle, stromal and immature endothelial cells at the single cell level (Figure 4H). Taken together, we describe novel culture conditions that allow for the maintenance and expansion of Gp38^+^PW1^+^ epicardial cells using hypoxic conditions and that these cells are multipotent progenitors for stromal cells, smooth muscle cells, and immature endothelial cells.

**Figure 4.**
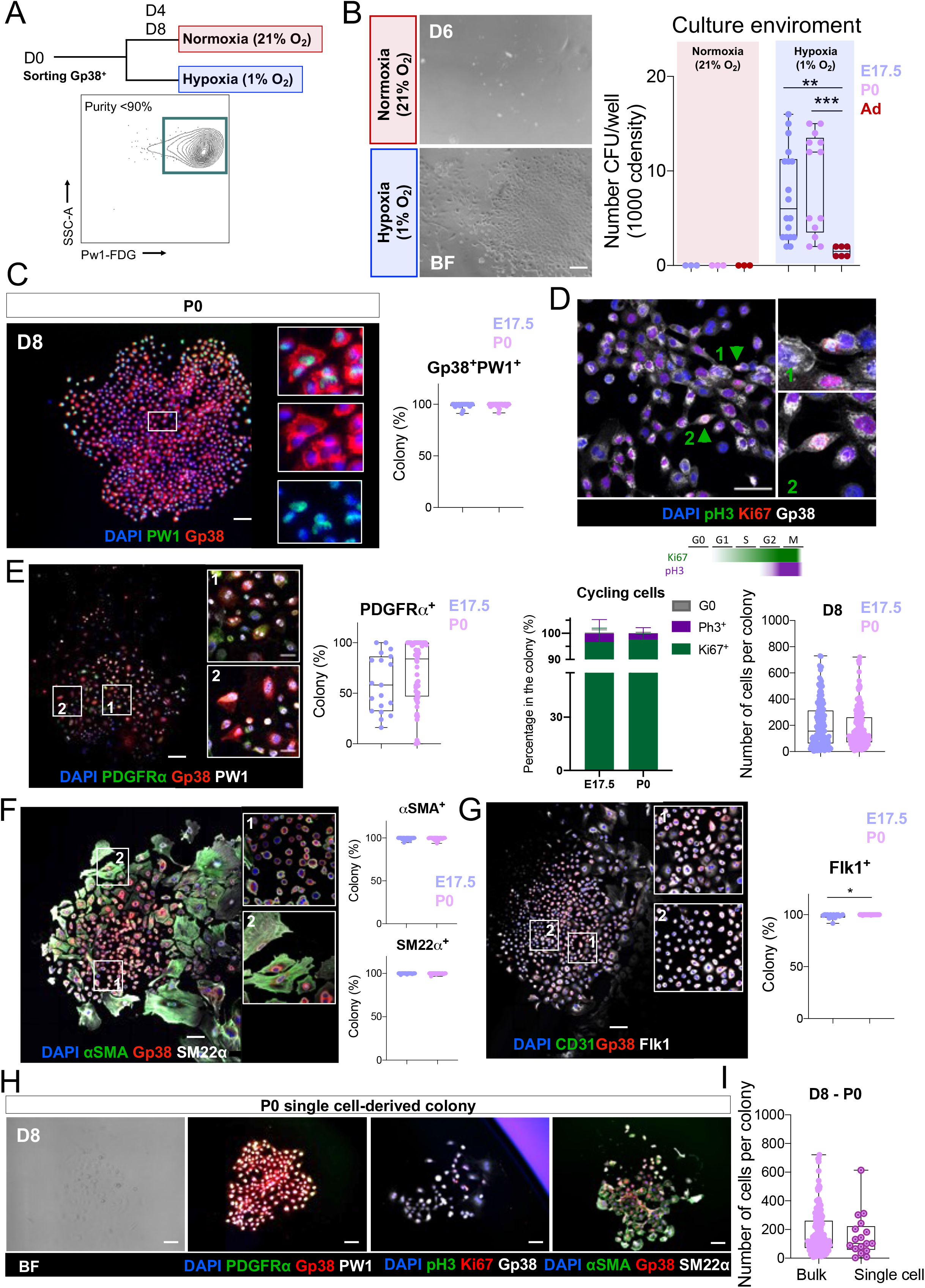
*In vitro* hypoxia maintains growth and cell fate potentials to Gp38^+^PW1^+^ epicardial cells. A. Schema of the *in vitro* experiment: cell sorting (Gp38^+^*Pw1*-FDG^+^ cells) with a purity of more than 90% from E17.5, P0, adult (Ad) and *in vivo* hypoxia-primed adult (H-Ad) hearts. Freshly sorted cells were cultured from D0 to D8 either in normoxia (21% O_2_) or hypoxia (1% O_2_). B. Frequency of colonies (1000 cells/well) cultured either under normoxia or hypoxia *in vitro* condition and the respective brightfield pictures of the cobblestone-like Gp38^+^ colonies. C. Immunolabeling shows the colonies keep the Gp38 and Pw1 expression similar to their initial *ex vivo* profile (clonogenicity). D. Immunolabeling of the cell cycle makers Ki67 and pH3 highlights the proliferative profile (green arrowheads) of colony of Gp38^+^PW1^+^ cells (self-renewal). E. Epicardial-derived and stromal cell markers (Gp38 and PDGFRα) are observed in the main cell fraction of the colony. F. Smooth muscle cell markers (αSMA and SM22α) are observed in the majority of the cells composing the Gp38^+^ colony. G. Endothelial marker Flk1, but not CD31, is observed to be expressed *in vitro* in Gp38^+^ colonies H. Single cell-derived colonies confirm the multipotential ability of the Gp38^+^PW1^+^ cells to differentiate into smooth muscle, stromal and immature endothelial cells. Scale bar: 100μm. All nuclei were counterstained with DAPI. Statistical significance was determined by Mann-Whitney test. \*\*\**p*<0.001, \*\**p*<0.01, \*\**p*<0.05. BF: Brightfield, E.17.5: embryonic day 17.5, P0: postnatal day 0, Ad: Adult.

### Physiological hypoxic priming induces the capacity to form CFU in adult epicardial cells

PW1^+^Gp38^+^ epicardial cells constitute a minor overall cell population in the adult heart (Figure 2A) and the epicardium is relatively quiescent(Cossette & Misra, 2011; Duim *et al.*, 2016; Manner *et al.*, 2005; Zhou *et al.*, 2008) and but hypoxic(Kocabas *et al.*, 2012). This raised the question whether *in vivo* hypoxia priming directly regulates epicardial progenitors. We compared the potential of PW1^+^Gp38^+^ epicardial cells derived from E17.5, P0, adult normoxia or adult hypoxia-primed mice. Limiting dilution analysis showed that P0 and E17.5 gave rise to epicardial colonies at similar frequency (1/25 and 1/40, respectively). Nevertheless, P0-derived PW1^+^Gp38^+^ epicardial cells showed a higher capacity to form colonies (CFUs) as compared to cells derived from E17.5 (Figure 5A), which may reflect that the epicardium is most active during early postnatal growth when the bulk of coronary vasculature is formed/matured(Tian *et al*, 2014). Moreover, adult epicardial cells from normoxic mice showed an 8-fold and 13-fold lower CFU capacity (1/324) as compared to fetal and newborn derived epicardial cells, respectively (Figure 5A). We therefore tested the capacity of PW1^+^Gp38^+^ epicardial cells derived from adult hearts following exposure to physiological hypoxia to test whether the level of oxygen accounted for CFU capacity. We found that the hypoxic ‘priming’ *in vivo* prior to culturing freshly isolated epicardial cells induced a 3-fold increase in the colony formation *in vitro* (Figure 5A-B). Furthermore, we found that the colonies derived from adult hypoxia-primed epicardial cells displayed a higher frequency of dividing cells (pH3^+^) as well as of PDGFRα^+^ cells similar to the behavior of colonies from newborn (P0) mice (Figure 5C-F), when the PW1^+^Gp38^+^ epicardial cells were more active. In conclusion, these observations demonstrate that hypoxia provides a critical stimulus for epicardial cell growth, expansion and differentiation and that the low oxygen levels exposure *in vivo* play a critical role in promoting these activities.

**Figure 5.**
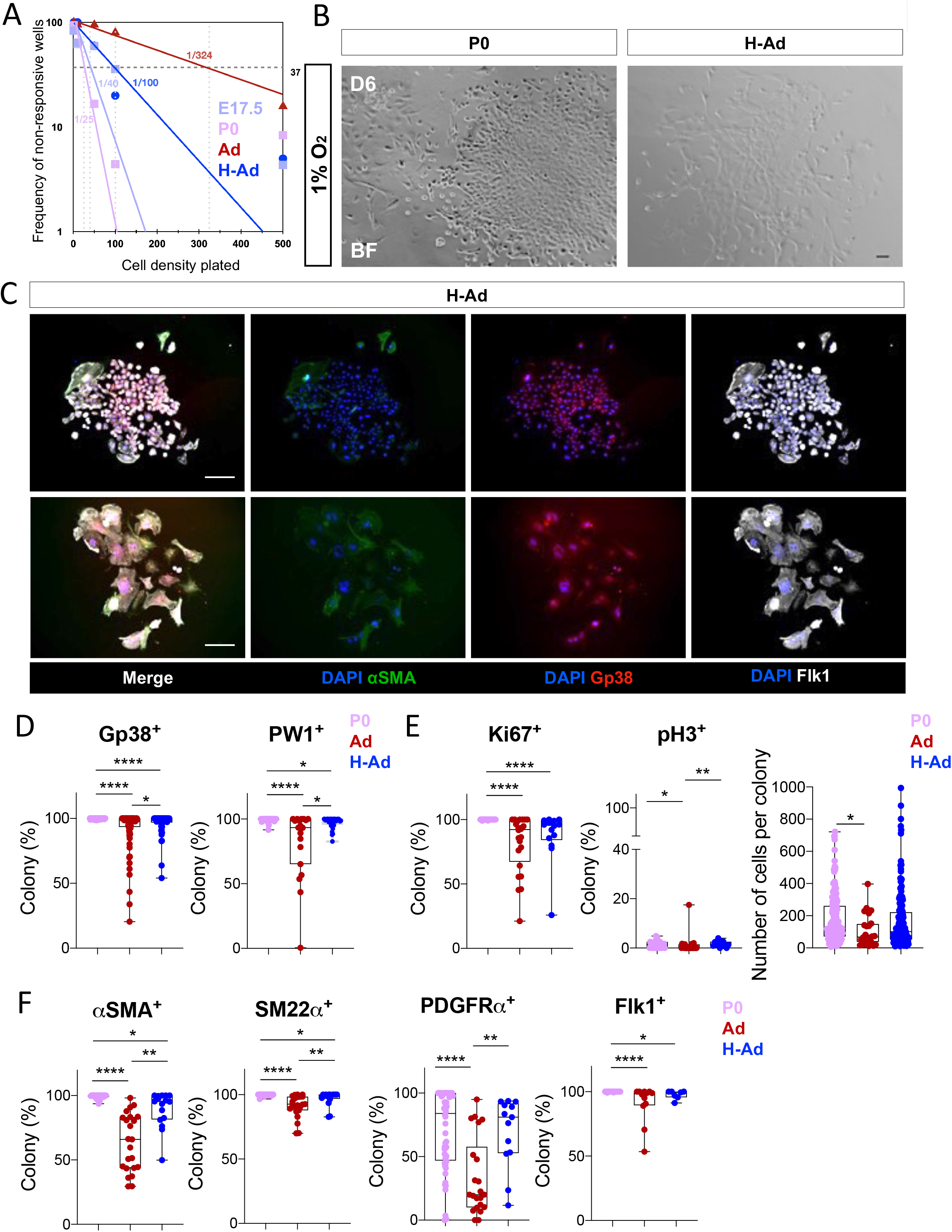
*In vivo* hypoxia-priming confers/restores the CFU ability in adult epicardial cells. A. Limiting dilution assay to determine (at the maximum likelihood parameter) the frequency of stem cells (CFU) within the Gp38^+^PW1^+^ cell population. The graph shows the number of negative wells (non-response) for the growth of a colony and the CFU probability (E17.5 - 1/40, P0 - 1/25, Ad - 1/324 and H-Ad - 1/100). B. Gp38^+^ cells form under hypoxia a cobblestone structure typical of epithelial structure colony (H-Ad), similarly to that observed in developing epicardium (E17.5 and P0). C. Immunolabelling of the hypoxia-primed adult Gp38^+^ colonies for relevant proteins of clonogenicity/self-renewal and multipotency. D. Comparison of the Gp38 and PW1 protein expression (clonogenicity/self-renewal) in P0, adult (normoxia and hypoxia). E. Comparison of the Ki67 and pH3 cell cycle protein (self-renewal) in P0, adult (normoxia and hypoxia). F. Comparison of the αSMA, SM22α, PDGFRα and Flk1 protein expression (multipotency) in P0, adult (normoxia and hypoxia). Scale bar: 100μm. All nuclei were counterstained with DAPI. The Maximum likelihood estimation (MLE) test (ELDA assay) was used for the statistical determination of the CFU values for each stage of Gp38^+^ cells with a *p*<0.0001. The line in the box plot represents the median. Statistical significance was determined by one-way ANOVA (Kruskal-Wallis test) with uncorrected Dunn’s test. \*\*\*\**p*<0.0001, \*\**p*<0.01, \**p*<0.05. P0: postnatal day 0, Ad: Adult, H: Hypoxia.

## Discussion

The epicardium is a mesothelial layer that surrounds the heart of all vertebrates and gives rise to multiple cell lineages during heart development and regeneration. There is considerable research effort aimed at identifying factors that activate the adult epicardium in order to promote a beneficial recovery following cardiac injury(Cao & Poss, 2018; Dube *et al*, 2017; Liu *et al*, 2014; Smart *et al.*, 2011; van Wijk *et al*, 2012; Zangi *et al*, 2013; Zangi *et al*, 2017; Zhou *et al.*, 2011; Zhou *et al*, 2012b). The embryonic and fetal hearts are physiologically hypoxic due to an underdeveloped cardiac vascular system and placenta restriction(Guimaraes-Camboa *et al.*, 2015; Nanka *et al.*, 2006; Olivetti *et al.*, 1980; Patterson & Zhang, 2010; Sayed *et al*, 2018; Tian *et al.*, 2013; Tian *et al.*, 2015; Tomanek, 1996; Wikenheiser *et al.*, 2006), whereas the hypoxic state is only maintained in the adult mammalian epicardium, as shown in this study and by others(Kocabas *et al.*, 2012). The epicardium has been shown to drive heart morphogenesis and serve as a reservoir of progenitors of non-cardiomyocyte lineages throughout life and in response to injury in non-mammalian vertebrates(Cai *et al.*, 2008; Dettman *et al.*, 1998; Gittenberger-de Groot *et al.*, 1998; Mikawa & Gourdie, 1996; Perez-Pomares *et al.*, 1997; Wessels *et al.*, 2012; Zhou *et al.*, 2008). The progenitor capacity of the adult mammalian epicardial cells is less explored, but is considered to be less robust. Additionally, whereas the epicardium is a source of coronary endothelial cells in non-mammalian vertebrates, it is not clear if this property is maintained in mammals.

We demonstrated previously that PW1 is broadly expressed during embryonic development and maintained in multiple adult progenitor niches(Besson *et al.*, 2011; Kuroiwa *et al.*, 1996; Mitchell *et al.*, 2010). We find a similar pattern of expression occurs in the heart with an initial widespread expression until early postnatal life and then restricted to the epicardium in the adult (Figure 6).

**Figure 6.**
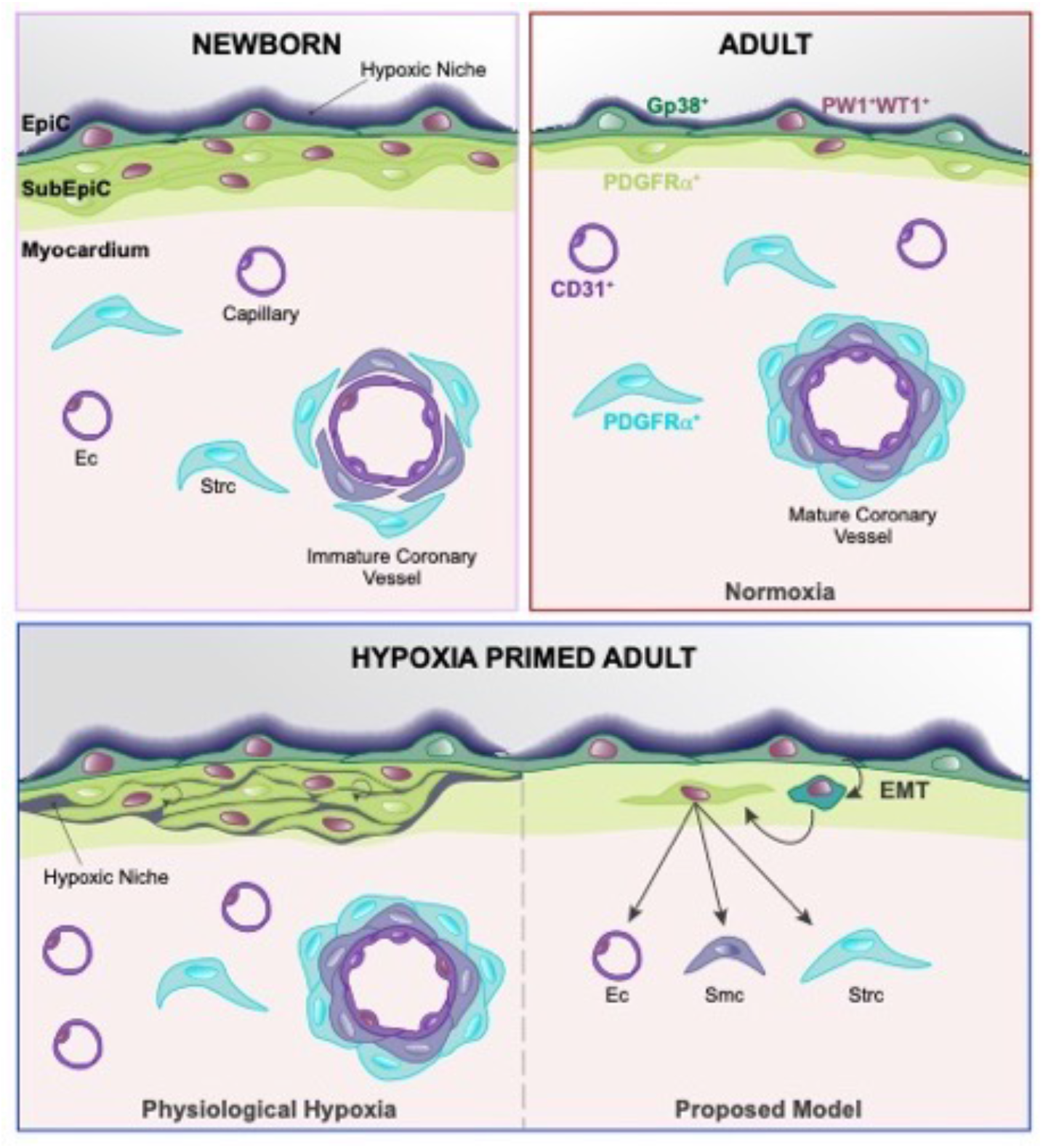
Proposed model for hypoxia activation of the adult murine epicardium to a progenitor cell function and multipotency. Epicardium is an active reservoir of progenitor cells that gives rise to coronary vasculature and stroma cells during perinatal life. The adult mammalian epicardium is a heart resident progenitor niche that maintains PW1 and WT1 expression, and a hypoxic environment. Hypoxia-primed epicardium activation induces a progenitor profile and shows multilineage differentiation ability, including the stromal, smooth muscle and endothelial lineage fates.

Although, definitive models for cell fate tracing are needed to clearly address whether hypoxia induces an endothelial cell fate in Gp38^+^PW1^+^ epicardial cells and our single cell transcriptional data provides a static picture of cells at the time of isolation, it nonetheless provides an unbiased approach that strongly supports a model in which the multipotent epicardium contributes to cells with both stromal and endothelial lineage profiles. Hypoxia has been shown to decrease fibrotic scarring coupled with functional recovery of the heart following ischemia(Nakada *et al.*, 2017) and we demonstrated previously that PW1 stromal cells contribute to fibrosis following myocardial infarction(Yaniz-Galende *et al.*, 2017). Here, our single cell transcriptome analysis shows that epicardial cells are a heterogenous population. Gp38^+^PW1^+^ epicardial cells show a strong lineage relationship between epicardial cells and PDGFRα^+^ cells (subepicardial and stromal cells) and, at less extent, but clearly more robust after hypoxia exposure, between epicardium and endothelial cells. The presence of a distinct subset of epicardial cells with endothelial lineage fate after hypoxia exposure is analogous to the developing epicardium profile, indicating the potential of adult resident epicardial cells to activate an immature endothelial cell program. We reported previously that cardiac PW1^+^ stromal cells can be purified based upon the expression of αV-integrin and that pharmacological inhibition of αV-integrin results in a potent inhibition of fibrosis in response to ischemia coupled with a reduced infarct size(Bouvet *et al.*, 2020). Whether hypoxia and αV-integrin action share a common signaling pathway endpoint will be of interest to pursue. These results provide a cellular basis by which physiological hypoxia promotes angiogenesis. The precise pathways acting in response to hypoxia likely involve well characterized cell stress pathways that are also regulated by *Pw1*, such as TNFα-NFκB signaling in cell growth and survival(Relaix *et al.*, 1998) and p53 signaling in apoptosis(Deng & Wu, 2000; Relaix *et al.*, 2000), as well as the glucose metabolism regulation(Correra *et al.*, 2018).

This study also provides a novel strategy to purify and culture Gp38^+^PW1^+^ epicardial cells that will prove invaluable to further dissect mechanisms underlying the role of epicardial cells during injury and repair. Perinatal-derived PW1^+^Gp38^+^ epicardial cells show clonogenicity, self-renewal and multipotency properties at the single cell level and can differentiate into PDGFRα^+^ cells (subepicardial and stromal cells), αSMA^+^SM22α^+^ smooth muscle cells and Flk1^+^ immature endothelial cells. This cell fate potential is dependent upon oxygen levels. We further show that epicardial cells isolated from the adult heart markedly improved following *in vivo* hypoxic priming their ability to be maintained and expanded *in vitro*. Taken together, this study demonstrates that resident Gp38^+^PW1^+^ epicardial cells retain progenitor competence in the adult mammalian heart and that hypoxia is critical to promote progenitor expansion and multi-lineage differentiation, including the activation of endothelial cell fates. These finding serve as a basis to design therapeutic approaches to promote heart revascularization and provide an invaluable *in vitro* model for further studies.

## Methods

A detailed methods section is provided as Online Supplement Material.

### Mice

C57BL/6J mice from Janvier and *PW1*^*IRESnLacZ*^ (*PW1*^*nlacZ*^) transgenic reporter mice(Besson *et al.*, 2011) were used. All animal procedures were approved by our institutional research committee (CEEA34 and French ministry of research) and followed the animal care guideline in Directive 2010/63/EU European Parliament.

### *In vivo* hypoxia

A cage connected to nitrogen and oxygen gas (ProOxP360, Biospherix) was used for the *in vivo* hypoxia with controlled atmosphere by silica gel (Carlo Erba) and soda lime (Intersurgical). Mice were exposed to hypoxia (10% O_2_) during 2 weeks.

### Cardiac cell suspension

Heart fragments were incubated at 37°C with enzymatic solution (Collagenase II and DNase I) for serial rounds of digestion.

### Flow cytometry and cell sorting

Cell suspensions were stained with a panel of surface antibodies, C_12_FDG to detect ß-galactosidase activity and viability dye before flow cytometry analysis. The list of antibodies for flow cytometry is provided in Supplemental Material.

### Primary cell culture and cell immunofluorescence staining

Sorted epicardial cells (Gp38^+^PW1^+^) were plated in amplification medium under normoxia (21 % O_2_) or acute hypoxia (1% O_2_) for 8 days. Cells were fixed, permeabilized and blocked. Primary antibodies were incubated followed by the adequate secondary antibodies and counterstained with DAPI. The list of antibodies for cell culture is provided in Supplemental Material.

### Histological processing and tissue immunofluorescence staining

Hearts were frozen, embedded in OCT and cut in four distinct coronal regions. Sections were fixed, permeabilized and blocked. Primary antibodies were incubated followed by the adequate secondary antibodies and counterstained with DAPI. The list of antibodies for histology is provided in Supplemental Material.

### X-gal staining

ß-galactosidase activity (LacZ) was detected in heart cryosections as described previously(Relaix *et al*, 2004; Sanes *et al*, 1986; Zakany *et al*, 1988).

### Pimonidazole injection and detection

Pimonidazole was injected intraperitonealy in adult mice (90 minutes incubation) and injected subcutaneously in newborn mice (3 hours incubation) before heart collection. Hearts were frozen and pimonidazole was detected with MAb1 (Hydroxyprobe), according to the manufacturer’s instructions.

### Multiplex qPCR (bulk and single cell)

Cells were sorted directly into RT-STA reaction mix (CellsDirect One-Step qRTPCR Kit) and 0.2x specific TaqMan Assay mix. Taqman assays are listed in Supplemental Material. Multiplex qPCR was preformed using the microfluidics Biomark HD system (Fluidigm) as previously described(Chea *et al*, 2016) for the same TaqMan Assay panel in Supplemental Material.

### Bioinformatic analysis

Gene expression raw data (Biomark, Fluidigm) of both bulk and single cell were normalized with Gapdh housekeeping gene, and ‘heatmap’, ‘Rphenograph’ and ‘diffusion map’ were created using R packages(Levine *et al*, 2015).

### Quantifications and statistical analyses

Histological images were analyzed with Icy software(de Chaumont *et al*, 2012). Graphs and statistics were performed with Prism software (GraphPad). The line in the box plots represents the median value. For the statistical comparison, two groups were tested by Mann-Whitney U test and for more than two groups one-way analysis of variance (ANOVA) was used. A value of p<0.05 was considered significant. Biological replicates represent independent experiments.

## Acknowledgements

We thank the members of D.S. and J.S.H. laboratories for fruitful discussions. We thank Ana Cumano (Institut Pasteur) and Zeina al Sayed (PARCC) for careful reading and discussion. We are grateful to the flow cytometry platforms (Catherine Blanc and Bénédicte Hoareau-Coudert from CyPS Pierre et Marie Curie University and to Camille Knosp from PARCC), to the animal facility platform for their help handling mouse lines (to the Pierre et Marie Curie University animal facility and to ERI at PARCC), to Sophie Nadaud for the technical help with the establishment of the *in vivo* hypoxia environment, to Priya Kantane for technical support, to the assistance in the Biomark (Valentina Libri and Valérie Feffer from Cytometry and Biomarkers UTechS, Institut Pasteur).

## Author contributions

A.S. and M.V. conceived and performed experiments, performed formal analysis, and wrote the manuscript; S.T. performed experiments and analysis; F.S.S. provided analysis assistance and discussion, G.M., D.S., J.S.H. provided feedback and reviewed the manuscript; D.S. and J.S.H. secured funding.

## Supplemental Materials

Expanded Methods

Supplemental Tables I

Supplemental

Figures I – VII

---

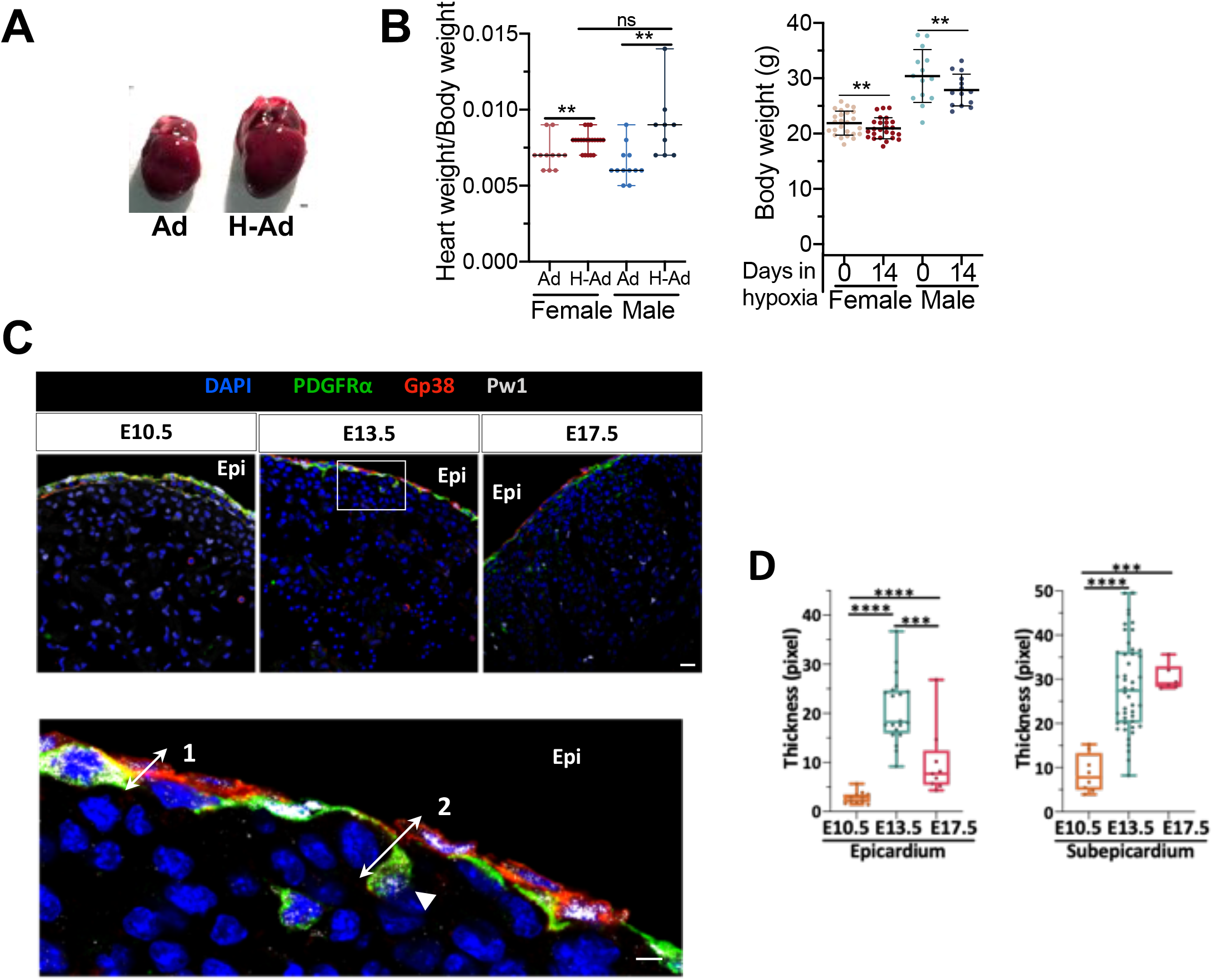

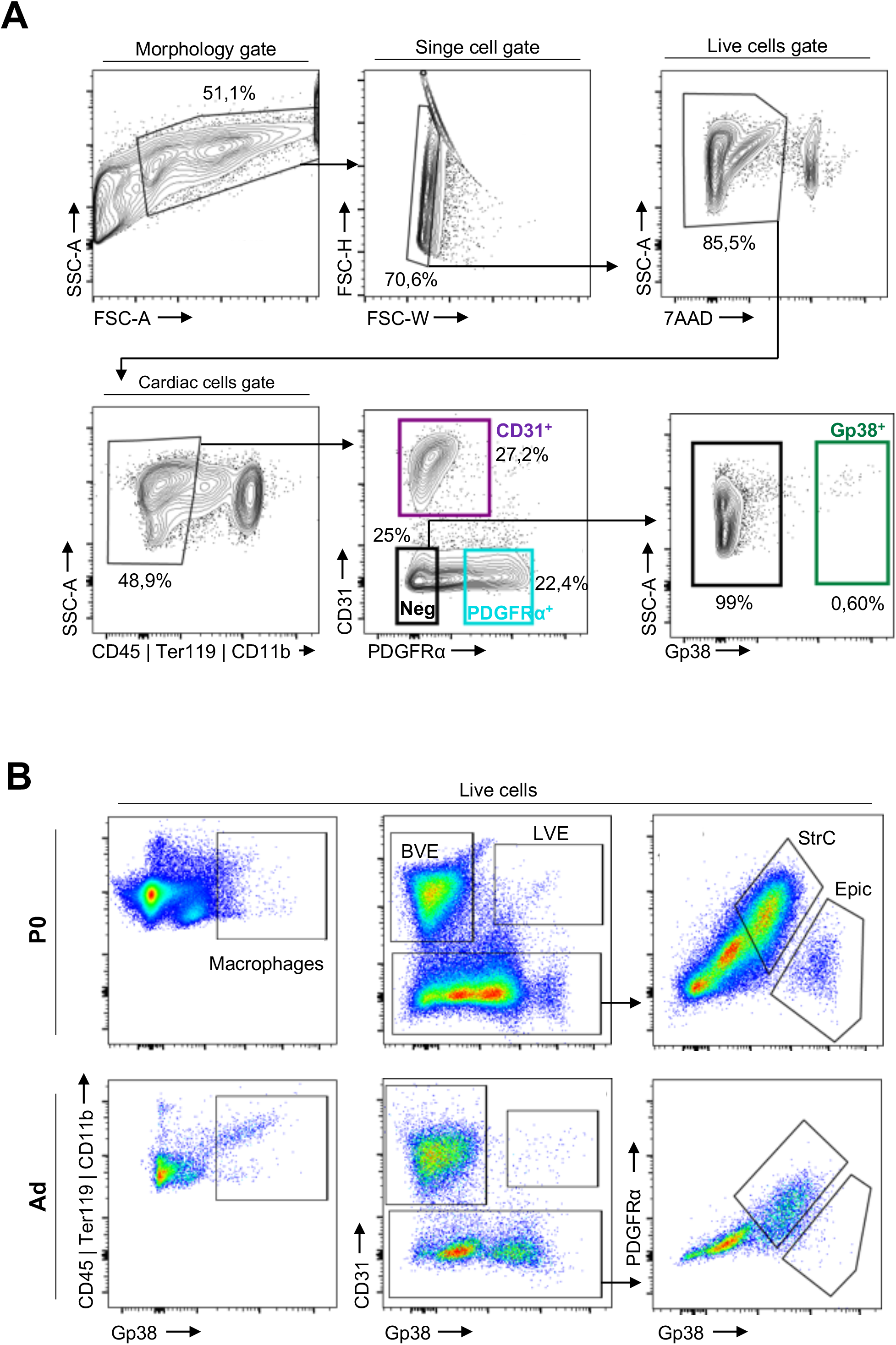

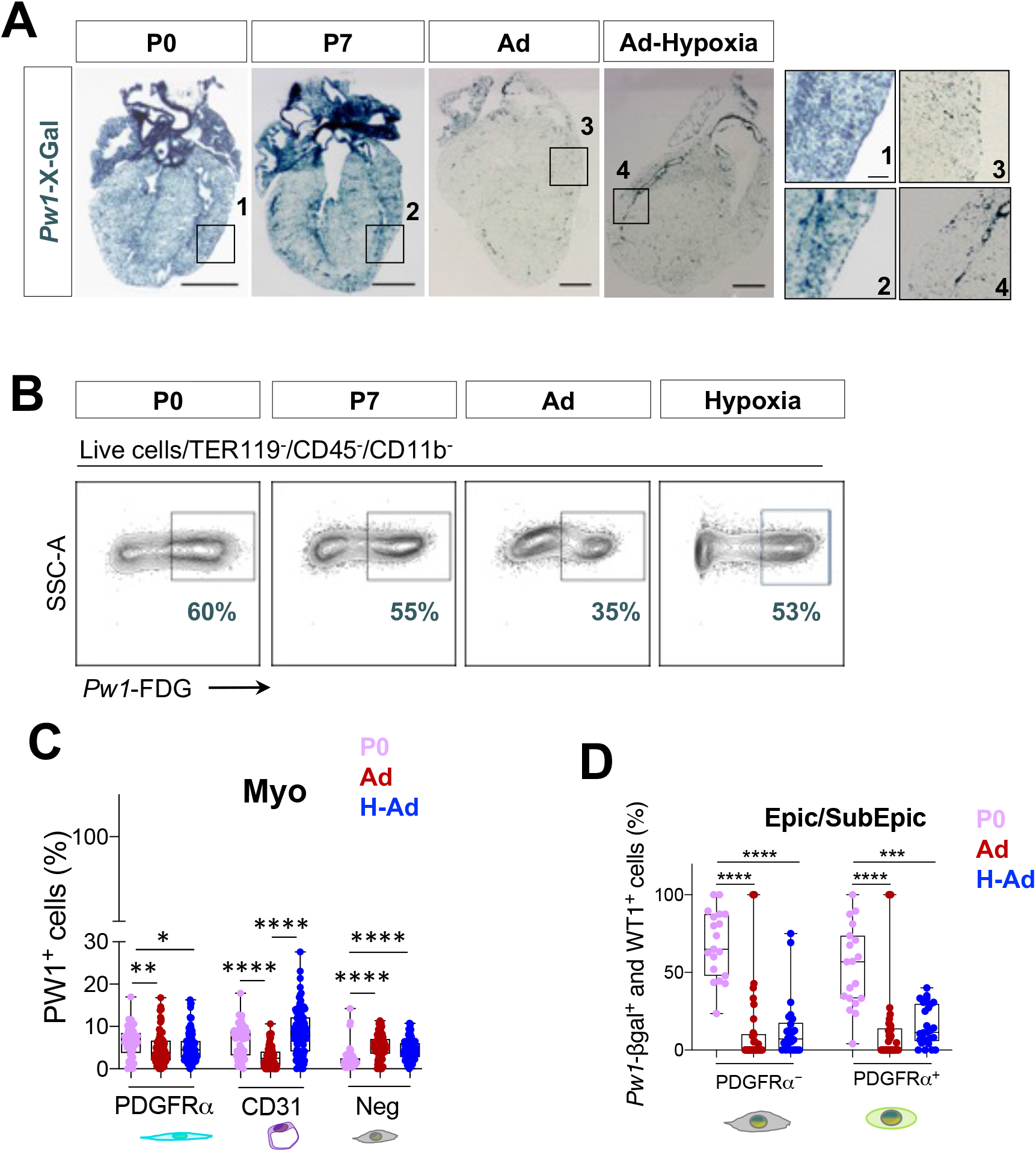

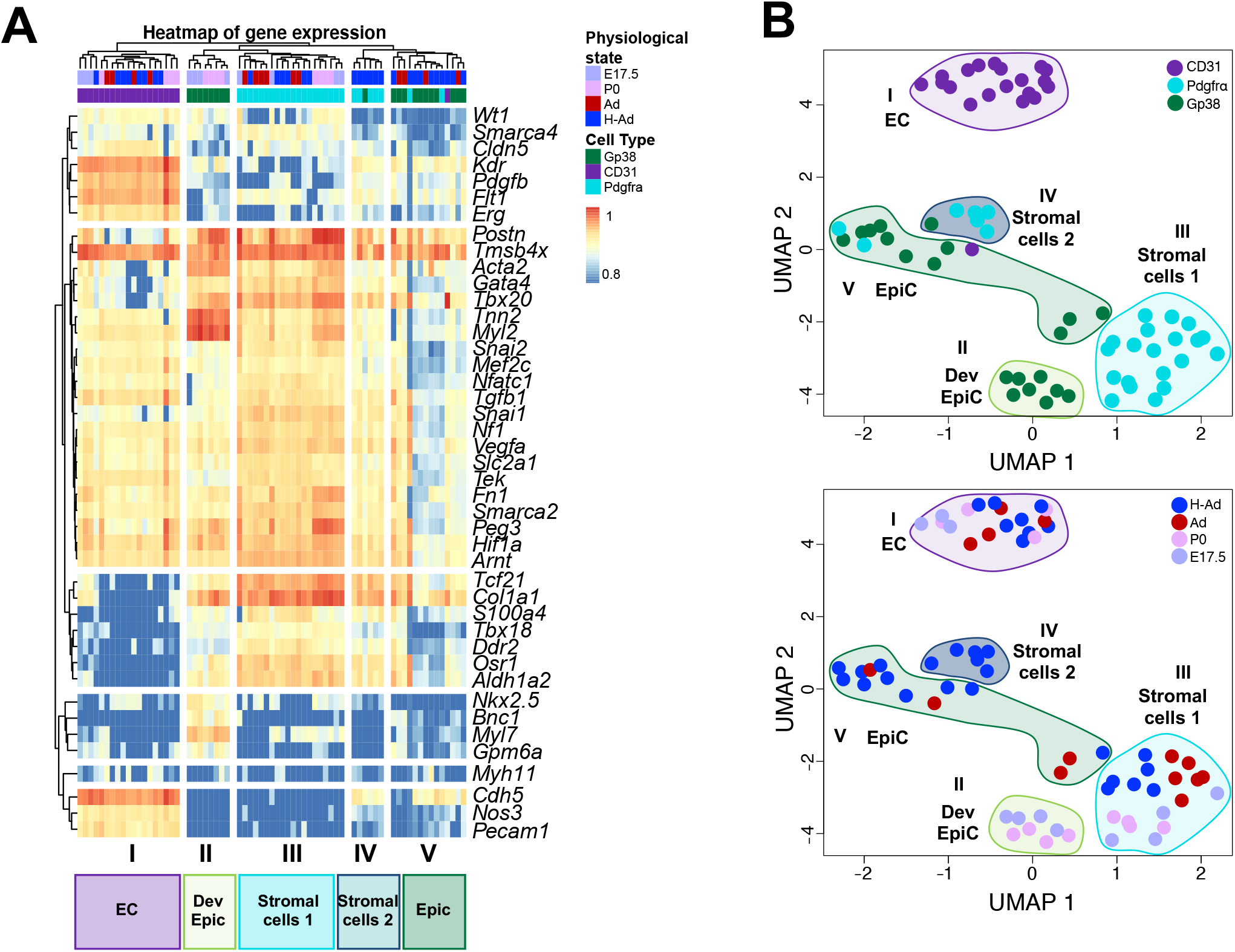

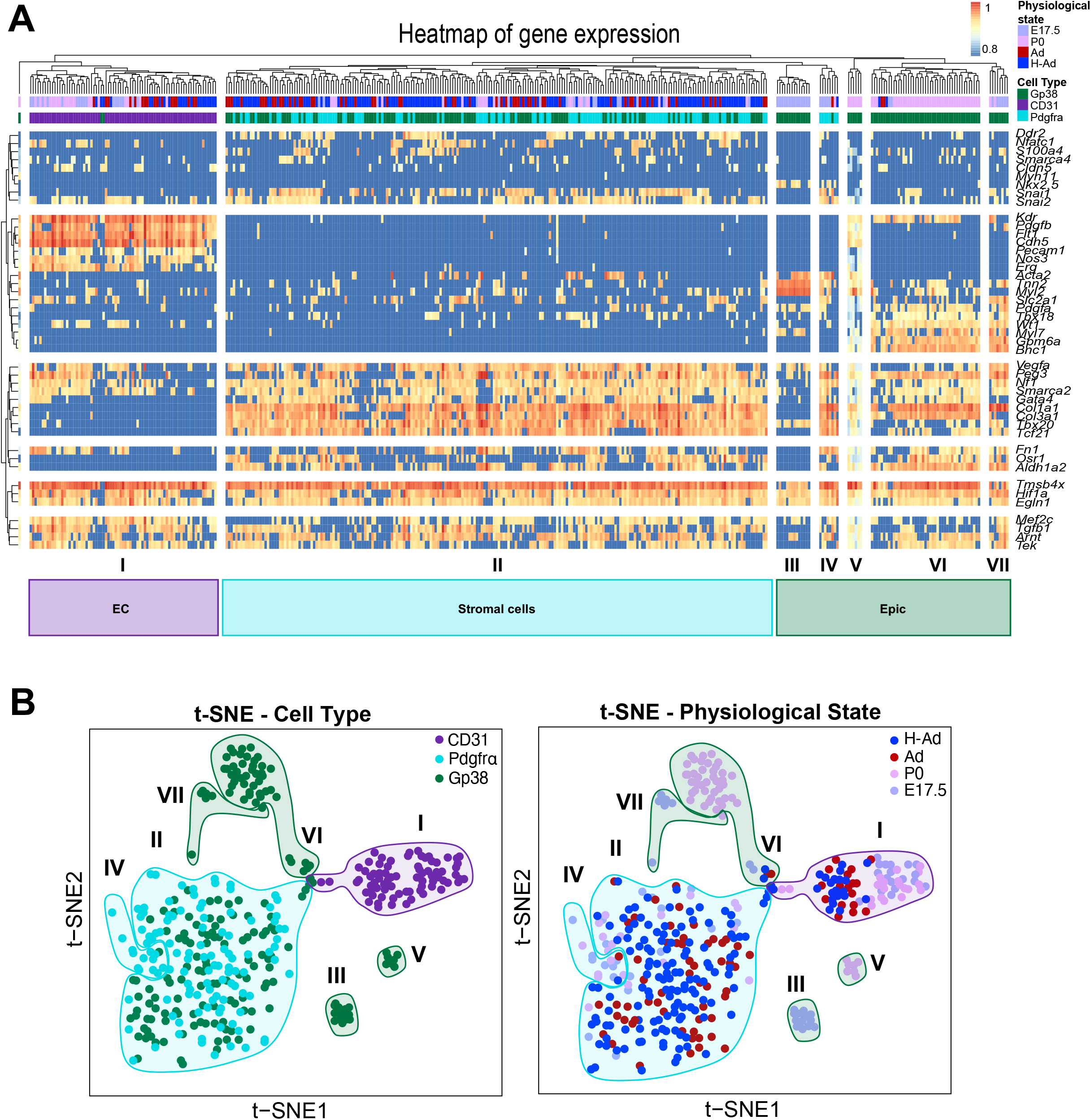

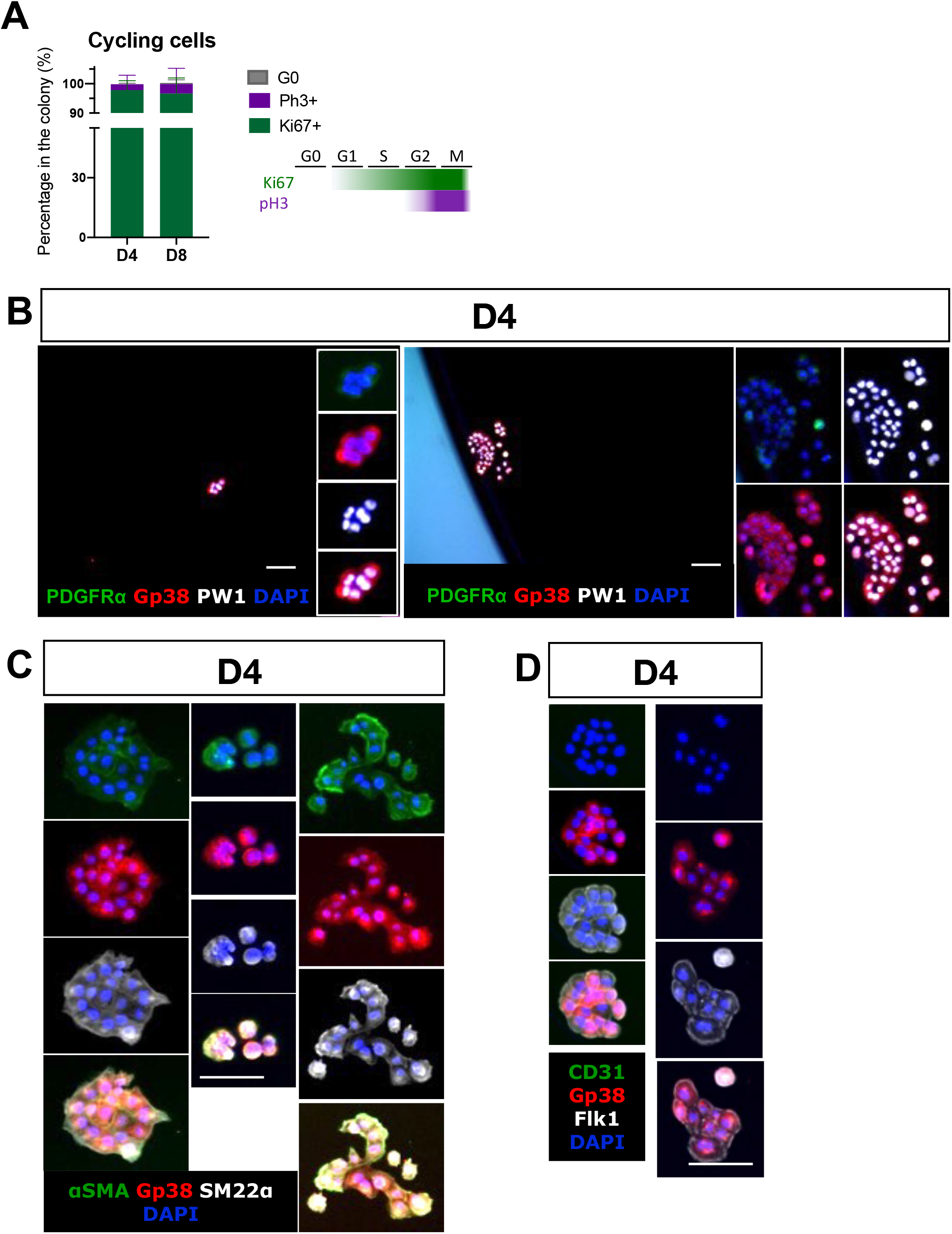

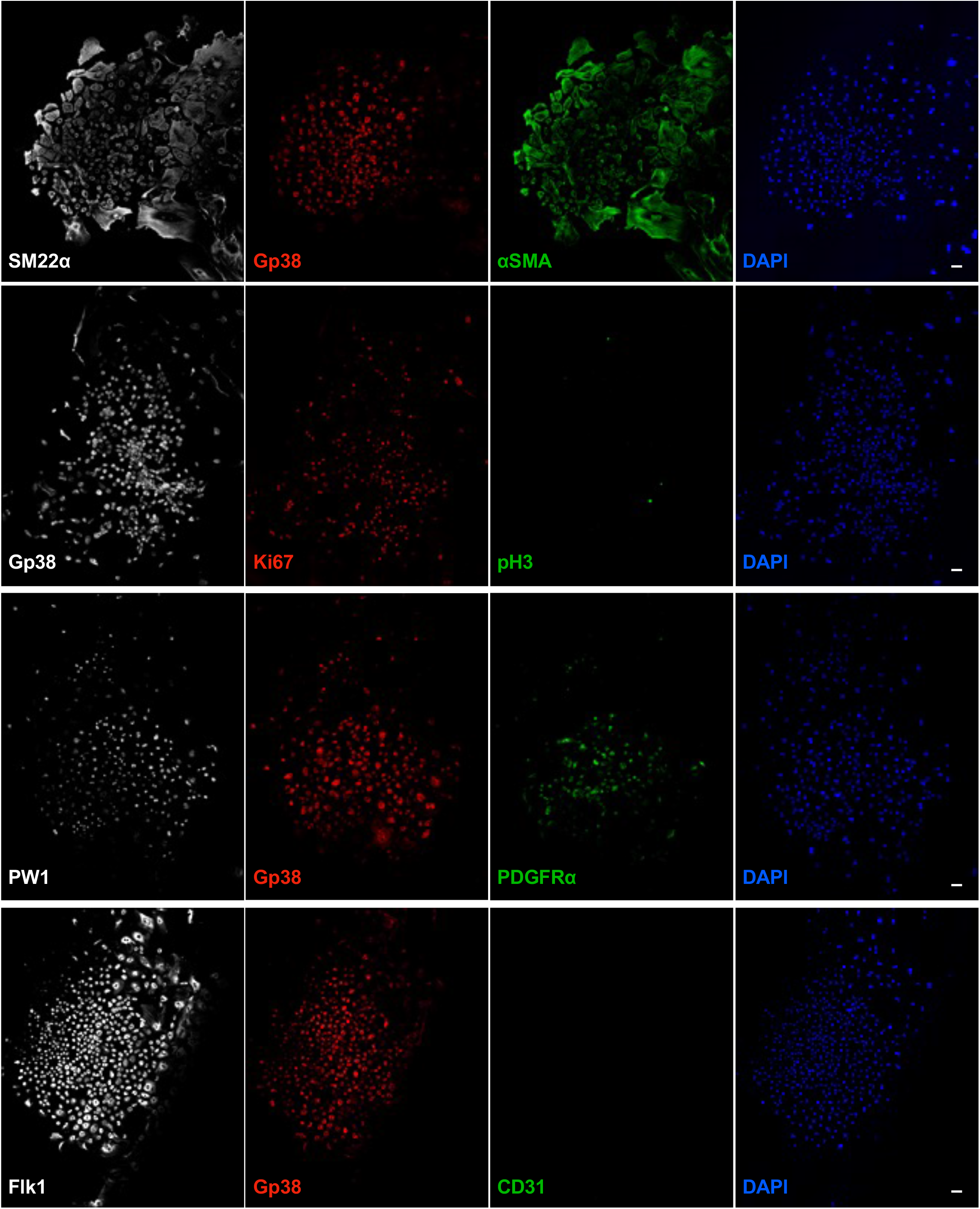

